# Natural scene and object perception based on statistical image features: psychophysics and EEG

**DOI:** 10.1101/2025.04.23.650126

**Authors:** Taiki Orima, Fumiya Kurosawa, Taisei Sekimoto, Isamu Motoyoshi

**Affiliations:** Department of Life Sciences, The University of Tokyo (153-8902, Komaba 3-8-1, Meguro-ku, Tokyo, Japan); Japan Society for the Promotion of Science; Center for Information and Neural Networks (CiNet), National Institute of Information and Communications Technology (NICT), Japan

**Author notes:** Author contribution: IM conceived, and TO, FK, TS, and IM designed the study. TO, FK, and TS conducted the experiments and analyzed the data. TO, FK, TS, and IM wrote the first draft of manuscript. TO and IM wrote the final version of manuscript. All authors approved the submitted version.

## Abstract

Recent studies have suggested the importance of statistical image features in both natural scene and object recognition, while the spatial layout or shape information is still important. In the present study, to investigate the roles of low- and high-level statistical image features in natural scene and object recognition, we conducted categorization tasks using a wide variety of natural scene and object images, along with two types of synthesized images: Portilla-Simoncelli (PS) synthesized images, which preserve low-level statistical features, and style-synthesized (SS) images, which retain higher-level statistical features. Behavioral experiments revealed that human observers (of either sex) could categorize style-synthesized versions of natural scene and object images with high accuracy. Furthermore, we recorded visual evoked potentials (VEPs) for the original, SS, and PS images and decoded natural scene and object categories using a support vector machine (SVM). Consistent with the behavioral results, natural scene categories were decoded with high accuracy within 200 ms after the stimulus onset. In contrast, object categories were successfully decoded only from VEPs for original images at later latencies. Finally, we examined whether style features could classify natural scene and object categories. The classification accuracy for natural scene categories showed a similar trend to the behavioral data, whereas that for object categories did not align with the behavioral results. Taken together, these findings suggest that although natural scene and object categories can be recognized relatively easily even when layout information is disrupted, the extent to which statistical features contribute to categorization differs between natural scenes and objects.

**Significance Statement:** Humans can reliably recognize complex natural scenes and objects. Recent studies have suggested that such recognition may rely on statistical image features, but the extent to which these features contribute to the recognition remains unclear. In the present study, we investigated how well statistical image features account for the perception of natural scenes and objects by conducting psychophysical categorization experiments and EEG decoding analyses. We found that natural scene categories could be reliably recognized based on statistical image features, and this recognition was consistent with neural responses. In contrast, although statistical image features also contributed to object category recognition, their effect appeared to be more limited. Together, these findings highlight the utility of statistical image features in visual perception.

## Introduction

Humans are capable of effortlessly and robustly recognizing natural scenes and objects (Potter, 1976; Wallis & Rolls, 1997; Quiroga et al., 2005; DiCarlo & Cox, 2007; DiCarlo et al., 2012; Epstein & Baker, 2019). Classical studies have emphasized that scene recognition relies on information beyond statistical image features, such as spatial layout (Biederman, 1972; Palmer, 1975; Davenport & Potter, 2004). The importance of information beyond statistical image features such as shape, contour, and global form has also been suggested in object recognition studies (Biederman, 1987; Biederman & Ju, 1988; Poggio & Edelman, 1990; Elder & Zucker, 1998).

On the other hand, previous psychophysical studies have demonstrated that humans can perceive the content of scenes and objects with brief presentation durations (Thorpe et al., 1996; VanRullen & Thorpe, 2001; Greene & Oliva, 2009). Such recognition of scenes and objects may rely on lower-level, relatively global, and more instantaneously processed features such as those computed by spatial envelope and scale-invariant feature transform (Oliva & Torralba, 2001; Lowe, 1999, 2004) than high-level information such as spatial layout, shape, and global form. Additionally, modern deep neural network (DNN) models that achieve remarkable object recognition often rely more heavily on texture-based information than on local shape features (Geirhos et al., 2018; Baker et al., 2018; Hermann et al., 2020; Li et al., 2020). Recent studies have even suggested that texture-based neural representations may exist in the human ventral visual pathway (Long et al., 2018; Ayzenberg & Behrmann, 2022; Jagadeesh & Gardner, 2022), which is responsible for object recognition (Goodale & Milner, 1992; Mishkin et al., 1983; Logothetis & Sheinberg, 1996; Ungerleider & Haxby, 1994; Kourtzi & Kanwisher, 2000; Grill-Spector et al., 2000; Grill-Spector et al., 2001; Cichy et al., 2014; Wurm & Caramazza, 2022). In fact, basic tasks such as animal detection can be performed with Portilla-Simoncelli (PS) synthesized natural scene images (Portilla & Simoncelli, 2000), which preserves statistical image features of the original image (Banno & Saiki, 2015).

Although the PS statistics alone seem to be still insufficient for scene and object recognition (Loschky et al., 2011), it remains possible that higher-level statistical image features contribute substantially to natural scene and object recognition. Recent advances in DNN-based object classification models (Krizhevsky et al., 2012; Simonyan & Zisserman, 2014) have enabled the extraction and utilization of such higher-order statistical image features as the style features (Gatys et al., 2015). As clearly demonstrated by neural style transfer (NST) applications using white noise as the content image (Orima et al., 2025), the style-synthesized images, which preserve style features of original images, capture the surface property of the original image. Therefore, the style features may serve as higher-level statistical image features, potentially contributing to the perception of natural scenes and objects.

To investigate this possibility, we examined human observers’ ability to categorize natural scene and object images into predefined 10 categories, using PS-synthesized images and style-synthesized images. Behavioral results revealed that for both natural scene and object images, observers could successfully categorize style-synthesized images. We further recorded visual evoked potentials (VEPs), and found that natural scene categories could be decoded from VEPs for original and style-synthesized images at statistically significant levels starting from 175 ms after the stimulus onset with high accuracy. In contrast, although object categories were also decoded from VEPs after 200 ms, the classification accuracy for style-synthesized images was only slightly above chance level. Finally, we revealed that natural scene categories could be predicted from style features in a manner consistent with behavioral data, whereas object categories showed little correspondence between classification based on style features and behavioral performance. These results suggest that higher-level statistical image features, style features, substantially contribute to the recognition of natural scene categories and are likely utilized by the human visual system. In contrast, style features also support object category recognition, but object recognition may rely on information that cannot be fully described by style features alone.

## 1: Psychophysical experiment for natural scene and object category classification

To examine the extent to which statistical image features contribute to natural scene and object recognition, we conducted behavioral experiments in which observers were asked to categorize either natural images or their corresponding Portilla-Simoncelli-synthesized (PS) and style-synthesized (SS) images that preserve lower-level and higher-level statistical image features of the original images, respectively.

### Materials & Methods

#### Apparatus (natural scene images)

For natural scene images, visual stimuli were generated using a PC (DELL PRECISION T1700), two HP EliteDesk 800 G5 Desktop Mini units, three HP ProDesk 400 G3 DM (Japan) units, one HP ProDesk 400 G3 Mini, one HP ProDesk 400 G6 Desktop Mini PC (Japan), and three HP Z2 Mini G4 Workstations. Visual stimuli were presented on LCD monitors, including three BENQ XL-2720-B, two BENQ XL-2730-Z, one BENQ XL-2731-K, two BENQ XL-2735-B, one BENQ XL-2746-K, one BENQ XL-2746-S, and one ZOWIE XL2746K. Due to COVID-19 restrictions, some observers completed the experiment in a dark room at home using their own monitors (same models as above). All monitors were gamma-corrected (ColorCal II, Cambridge Research Systems), and the refresh rate was set to 60 Hz. The mean background luminance of the monitors ranged from 38 to 55 cd/m². Viewing distances were adjusted so that the spatial resolution was 0.022 degrees per pixel.

#### Apparatus (object images)

For object images, three observers completed the task in a dark room at the laboratory, viewing visual stimuli presented on an LCD monitor (BenQ XL2730B). The mean background luminance was 38.4 cd/m². To accommodate COVID-19 restrictions, eight additional observers completed the experiment in a dark room at home, using one of the following monitors: two BENQ XL2720B, two BENQ XL2730Z, one BENQ XL2735B, one BENQ XL2746S, one BENQ XL2731K, or one BENQ XL2746K. The mean background luminance ranged from 39 to 55 cd/m². All monitors were gamma-corrected (ColorCal II, Cambridge Research Systems), and the refresh rate was set to 60 Hz. Viewing distances were adjusted so that the spatial resolution was 0.022 degrees per pixel.

#### Observers (natural scene images)

Three authors and eight naïve students (ages 21–30, mean age = 23.8 years) participated in the experiment. All observers had normal or corrected-to-normal vision. The present study was approved by the “Ethical Review Committee for Experimental Research on Human Subjects, Faculty of Arts and Sciences, The University of Tokyo.” Written informed consent was obtained from all observers prior to the experiment.

#### Observers (object images)

A total of eleven observers took part in the experiment, including three authors and eight naïve students (ages 21–27, mean age = 23.5 years). All observers had normal or corrected-to-normal vision. The experiment was approved by the “Ethical Review Committee for Experimental Research on Human Subjects, Faculty of Arts and Sciences, The University of Tokyo.” Written informed consent was obtained from all observers.

#### Visual stimulus (natural scene images)

The visual stimuli consisted of 1,250 images in total, including 250 natural images (5.7 deg × 5.7 deg), their corresponding PS-synthesized images, and three types of style-synthesized images (SS3, SS4, and SS5) (Figure 1a). The natural scene images used in the present study were collected from publicly available online databases, including the SUN database (Xiao et al., 2010) and Places365 (Zhou et al., 2014). Natural scene images were selected based on criteria commonly used in previous studies (Epstein & Kanwisher, 1998; Oliva & Torralba, 2001; Greene & Oliva, 2009; Groen et al., 2013; Greene & Hansen, 2020, etc.), such that no single, specific object dominated the majority of the image, and the depicted scene could be identified based on its semantic structure and spatial layout. These images were assumed to have been captured with gamma 2.0 and loaded with gamma 0.5. All natural scene images were categorized into one of ten natural scene categories: bedroom, coast, forest, highway, inside city, kitchen, living room, mountain, office, and tall building.

**Figure 1.**
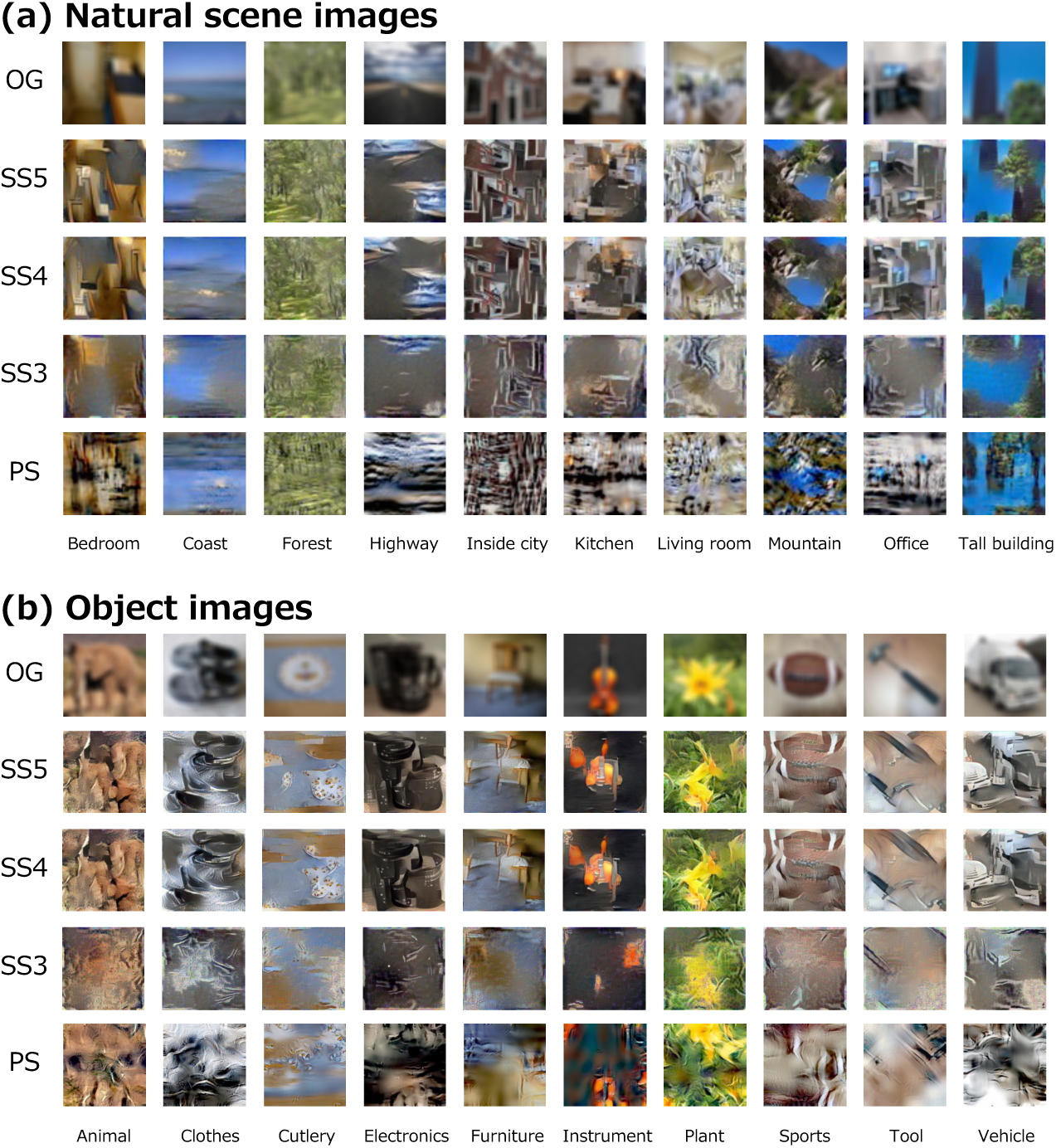
(a) Example images of natural scenes used in the experiment. OG indicates the original natural scene images, with one representative image shown for each of the ten categories: bedroom, coast, forest, highway, inside city, kitchen, living room, mountain, office, and tall building (from left to right). PS indicates PS-synthesized images, and SS3, SS4, and SS5 indicate three types of style-synthesized images. The PS and style-synthesized images shown here were generated from the corresponding OG images. (b) Example images of objects used in the experiment. The format is the same as in (a), with images arranged in the same order. We note that natural images were blurred to address copyright issues.

The PS-synthesized images were generated using the texture synthesis algorithm developed by Portilla & Simoncelli (2000), with the following parameters: number of scales of 4, number of orientations of 4, number of positions of 7, and number of iterations of 50. These parameters are the default values for PS-synthesis algorithm except for the number of iterations, and were adopted based on the results validated in our previous studies as well as other related study (Okazawa et al., 2015; Orima & Motoyoshi, 2021). In this PS-synthesis algorithm, the generated image retains only the moment statistics of the luminance image, the moment statistics of spatial frequency and orientation subbands, and the correlations between subbands. Previous studies have suggested that such statistical features are computed and processed in early visual areas such as V1 and V2, and contribute to texture perception (DeAngelis et al., 1999; Ferster & Miller, 2000; Bredfelt & Ringach, 2002; Freeman & Simoncelli, 2011; Freeman et al., 2012; Ziemba et al., 2019; Henderson et al., 2023).

Style-synthesized images were generated using Neural Style Transfer (NST) based on the method proposed by Gatys et al. (2015). NST enables the transfer of an artistic style from one image onto the content of another by extracting content information (such as contours and shapes) and style information (such as surface texture patterns) using a deep convolutional neural network (dCNN), and optimizing a loss function combining both content and style features. In the present study, using pre-trained VGG19, the style loss was computed from the original natural images, while the content loss was computed from white noise images, enabling us to generate style-synthesized images that retain only the style information of the original image but not its structural content. In detail, to manipulate the amount of style information extracted, we varied the convolutional layers of pre-trained VGG19 used for computing the style loss, generating three types of style-synthesized images. Specifically, style loss was computed from: conv1_1, conv2_1, and conv3_1 for SS3 images, conv1_1, conv2_1, conv3_1, and conv4_1 for SS4 images, and conv1_1, conv2_1, conv3_1, conv4_1, and conv5_1 for SS5 images. Based on the fact that shallower layers of dCNN models encode low-level image features such as spatial frequency and orientation statistics, and deeper layers of dCNN models do higher-level image features such as specific objects (Zeiler & Fergus, 2014; Kriegeskorte, 2015), SS3 images, which retains only shallow layers’ style features, were thought to share merely low-level image features with the original images, while SS4 and SS5 images, which retains deeper layers’ style features, were assumed to have higher-level image features with the original images. These style features were also utilized in the previous study and regarded as statistical image features (Jagadeesh & Gardner, 2022). As confirmed in another previous study (Orima et al., 2025), neural style transfer enables us to extract material property such as glossiness of surfaces, which surpasses what low-level statistical image features such as PS-statistics can describe.

#### Visual stimulus (object images)

The visual stimuli consisted of 1,000 images in total, including 200 original object images (5.7 deg × 5.7 deg) and their corresponding synthesized images (PS, SS3, SS4, and SS5) (Figure 1b). The synthesized images were generated based on the OG images using the same procedures as those applied to the natural scene images.

The object images were collected from online sources. In contrast to natural scene images, object images were selected such that a single object prominently occupied the central region of the image. Images depicting multiple objects or objects with unclear contours, which might hinder the perception of a single, isolated object, were excluded from the experiment. The images were assumed to have been captured with gamma 2.0 and loaded with gamma 0.5.

All object images were categorized into one of ten object categories: animal, clothes, cutlery, electronics, furniture, instrument, plant, sports, tool, and vehicle.

#### Experimental design and statistical analyses

##### Validation of four types of synthesized images

To verify whether the stimulus manipulations for each type of synthesized image (PS, SS3, SS4, and SS5) were implemented as intended, we conducted a perceptual evaluation experiment. The PS images preserved relatively low-level image statistics, referred to as Portilla-Simoncelli statistics, while the SSx images shared statistical properties with the original images as encoded in the X-th convolutional block of VGG19. According to previous studies (e.g., Zeiler & Fergus, 2014), SS3 is expected to retain image features that are nearly equivalent to those in PS, whereas SS4 and SS5 are assumed to preserve higher-level image features not captured in PS or SS3. Therefore, it is hypothesized that SS4 and SS5 should be perceptually more similar to the original images (OG) than PS or SS3.

To test this hypothesis, we conducted a ranking task, and seven observers (excluding the authors) participated in the experiment. For each natural scene or object image, the observers were asked to rank the synthesized images (PS, SS3, SS4, and SS5) in order of perceived similarity to the corresponding original image (OG). In each trial, the OG image was presented on the far left of the screen, while the four corresponding synthesized images were shown to its right in random order. Participants were allowed to move their eyes freely and instructed to rank the images by pressing keys in order of decreasing similarity to the OG (e.g., the button of closest synthesized image to the OG was pressed first).

For the analysis, we recorded the similarity ranking assigned to each synthesized image for every original image, averaged the ranks across images for each observer to obtain representative values, and then computed the means across observers. Statistical significance was evaluated using a one-sample t-test with the number of participants (*N* = 7) as the sample size.

##### Behavioral categorization of PS- and style-synthesized images

For natural scene images, observers viewed each of the 1,250 images once at the center of their visual field and were asked to categorize the presented image into one of ten predefined natural scene categories: bedroom, coast, forest, highway, inside city, kitchen, living room, mountain, office, or tall building. The order of image presentation was randomized across all trials: all versions of images were presented in random order. The experiment consisted of 25 short sessions, each comprising 50 trials. In each trial, a visual stimulus was presented for 200 ms at the fixation point. After stimulus presentation, observers selected the category of the image by referring to a list of the ten natural scene categories and responding via button press. Once a response was made, the trial ended, and observers initiated the next trial by pressing a button.

For object images, observers viewed each of the 1,000 object images once at the center of their visual field. For each image, they were asked to categorize the presented image into one of ten predefined object categories: animal, clothes, cutlery, electronics, furniture, instrument, plant, sports, tool, and vehicle. Observers were instructed to choose the category that best matched their perceptual impression of the presented image. They were explicitly told not to respond based on a small part of the image or to guess based on previous trials. When unsure of the correct category, observers were encouraged to choose the one that seemed most appropriate. The order of image presentation was randomized across trials, regardless of image type (OG, PS, SS3, SS4, SS5). In each trial, a stimulus was presented for 200 ms at the fixation point. After viewing the stimulus, observers selected the category by referring to the list of ten options and responded via button press. Each trial ended upon the observer’s response, and the next trial began when the observer pressed a button to proceed. No time limit was imposed for responses. Before the main experiment, all observers completed a short practice session consisting of 50 trials using a separate set of images not included in the main experiment, to familiarize themselves with the task.

The significance of behavioral data was tested by t-test using the number of images (250 for natural scene, 200 for object) as the sample size. We chose the number of images as the sample size because we sought to preserve generalizability across images (c.f. Yarkoni, 2022). The number of images were determined at the beginning of the whole experiment, and it never be changed. We applied the false discovery rate (FDR) correction method proposed by Benjamini & Hochberg (1995) to the p-values calculated at each data point, to address the multiple comparisons.

### Results

#### Sanity of four types of synthesized images

Figure 2 illustrates the results of the perceptual similarity judgments between the original images (OG) and the synthesized images. The psychophysical experiment revealed that, on average, SS5 was perceived as the most similar to the OG, followed by SS4, for both natural scene and object images. Furthermore, both PS and SS3 were rated as significantly less similar to the OG compared to SS4 (|*t*(6)|=17.7, *p*=2.1×10^-6^ for natural scene images; |*t*(6)|=23.4, *p*=4.0×10^-7^ for object images). These results are consistent with our hypothesis, indicating that SS4 and SS5 retain higher-level image features beyond those preserved in PS and SS3.

**Figure 2.**
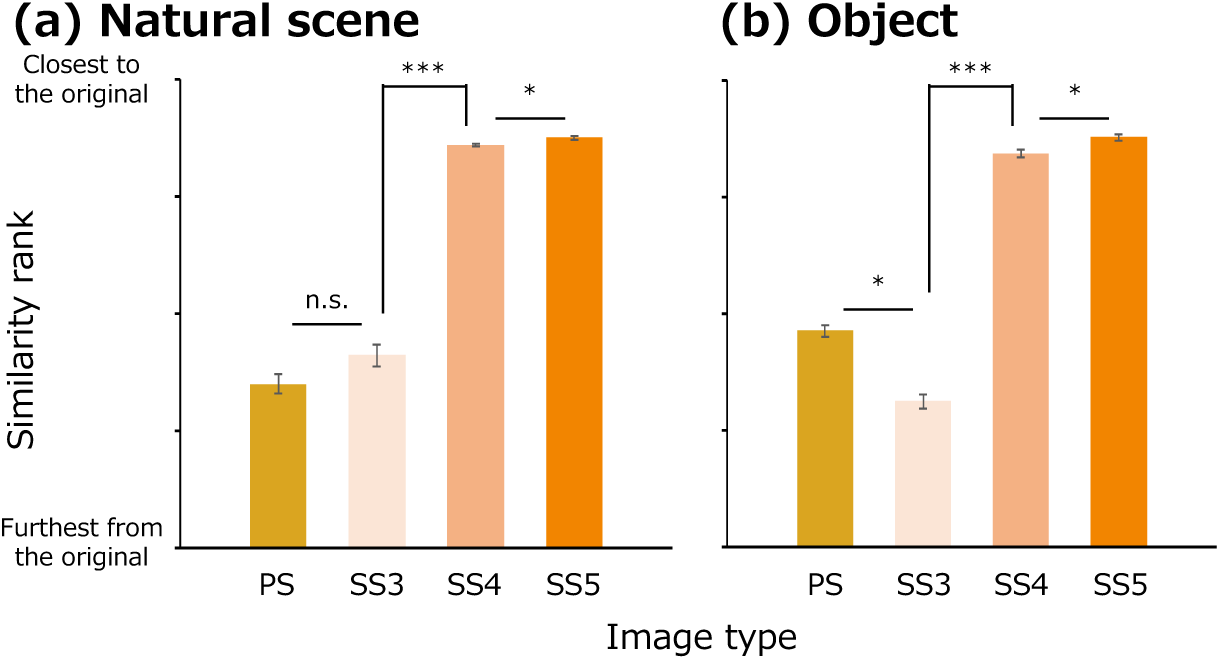
(a) Similarity rank of PS, SS3, SS4, and SS5 images in comparison to corresponding OG natural scene images. The higher value indicates synthesized image was closer to the original image. ***, *, n.s. shows the difference between the two conditions were statistically significant (p<0.001, p<0.05; FDR-corrected), and not statistically significant, respectively. (b) Similarity rank of PS, SS3, SS4, and SS5 images in comparison to corresponding OG object images. The format is the same as in (a).

#### Categorization accuracy across image synthesis conditions for natural scene and object images

Figure 3a shows the mean accuracy of observers in categorizing natural scene images across different synthesis conditions. The average categorization accuracy for style-synthesized images using three convolutional layers (SS3) was 30 %, which was lower than the 41 % accuracy for PS-synthesized images (PS). In contrast, style-synthesized images that incorporated a greater number of convolutional layers exhibited substantially higher performance: 71 % for SS4 and 73 % for SS5. Although these scores were far above those for PS images, they remained lower than those for original images (OG), which reached an average accuracy of 98 %. Two-tailed t-tests were performed using the number of images (n = 250) as the sample size. Categorization accuracy under all conditions was significantly higher than the chance level (10 %) for natural scene categorization (PS: *t*(249) = 15.9, *p* < 10^-38^; SS3: *t*(249) = 10.5, *p* < 10^-20^; SS4: *t*(249) = 35.9, *p* < 10^-99^; SS5: *t*(249) = 37.4, *p* < 10^-103^; OG: *t*(249) = 244.0, *p* < 10^-297^).

**Figure 3.**
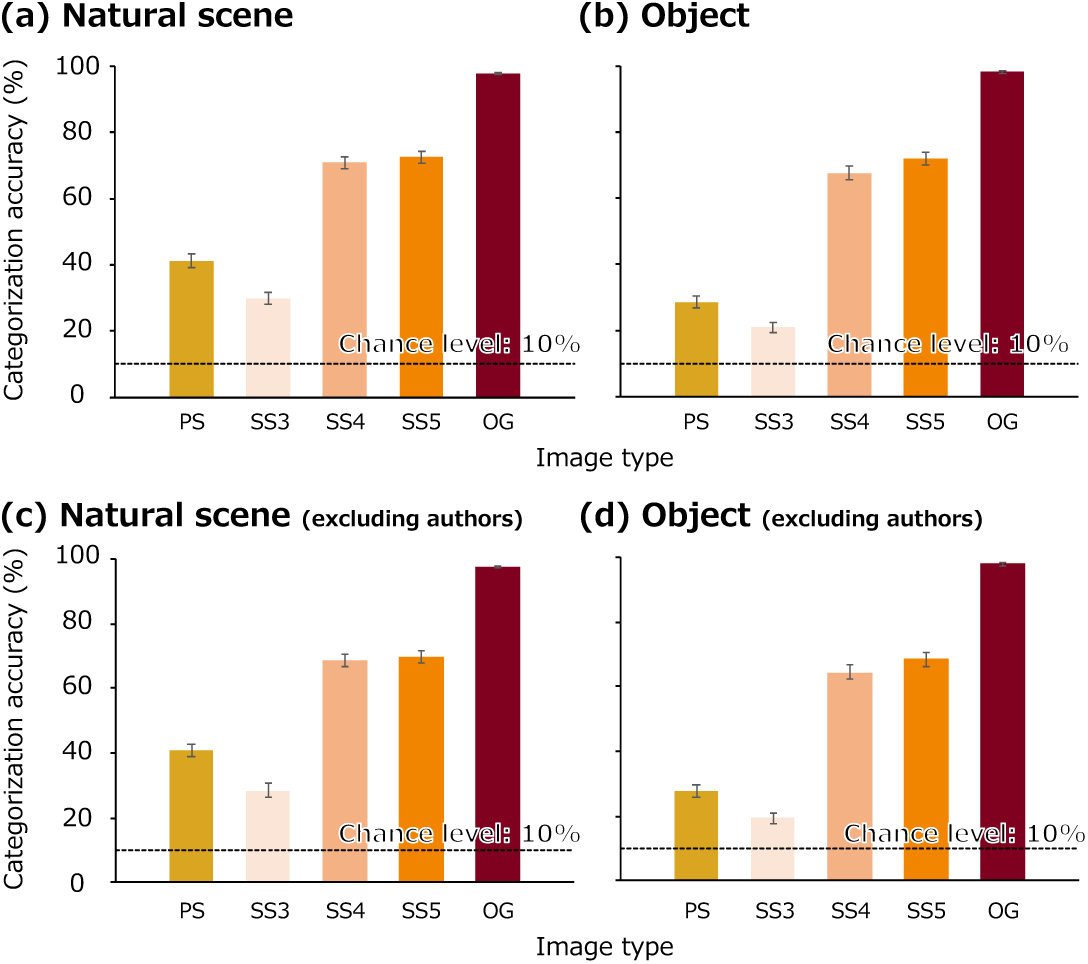
(a) Mean accuracy in the natural scene categorization task. The yellow bar represents the PS-synthesized images (PS), the orange bars represent the three types of style-synthesized images (SS3, SS4, SS5), and the red bar represents the original images (OG). The dotted line indicates the chance level (10 %) for this task. Error bars represent ±1 standard error of the mean (SEM) across images. (b) Results of the object categorization task. (c) Results of the natural scene categorization task excluding authors’ data. (d) Results of the object categorization task excluding authors’ data. Each bar shows the mean categorization accuracy for object images under each condition. The yellow bar corresponds to the PS-synthesized images (PS), the orange bars correspond to the style-synthesized images (SS3, SS4, SS5 from left to right), and the red bar corresponds to the original images (OG). The dotted line indicates the chance level (10 %). Error bars indicate ±1 SEM across images.

Figure 3b shows the mean categorization accuracy for object category judgments. In style-synthesized conditions (SS4 and SS5), object categories were identified more accurately than in PS or SS3 conditions, reaching approximately 70 % accuracy, which was comparable with the accuracy observed for original images (OG), which exceeded 98 %. Two-tailed t-tests (n = 200) confirmed that categorization accuracy under all synthesis conditions was significantly above the chance level (10 %) (PS: *t*(199) = 10.2, *p* < 10^-19^; SS3: *t*(199) = 7.27, *p* < 10^-11^; SS4: *t*(199) = 27.5, *p* < 10^-69^; SS5: *t*(199) = 31.9, *p* < 10^-79^; OG: *t*(199) = 241.9, *p* < 10^-246^). Figure 3c and 3d shows that these results remained significant both for natural scene category (PS: *t*(249) = 19.7, *p* < 10^-51^; SS3: *t*(249) = 14.7, *p* < 10^-35^; SS4: *t*(249) = 36.5, *p* < 10^-101^; SS5: *t*(249) = 38.2, *p* < 10^-105^; OG: *t*(249) = 225.3, *p* < 10^-289^) and for object category (PS: *t*(199) = 14.9, *p* < 10^-33^; SS3: *t*(199) = 12.5, *p* < 10^-26^; SS4: *t*(199) = 29.1, *p* < 10^-72^; SS5: *t*(199) = 32.9, *p* < 10^-81^; OG: *t*(199) = 231.6, *p* < 10^-243^) when the behavioral data form authors, who were familiar with the task, were excluded.

#### Categorization accuracy for each natural scene and object category

Figure 4a shows the mean accuracy of natural scene categorization for each category. For the original images (OG), accuracy was consistently high across all categories. Regarding the synthesized images, although the overall trend showed higher categorization accuracy for SS4 and SS5 compared to PS and SS3 images, the extent of this improvement varied across categories. For example, the bedroom category showed a substantial gap in accuracy between PS/SS3 and SS4/SS5 conditions, whereas in the coast and forest categories, the differences between PS/SS3 and SS4/SS5 were relatively small. Two-tailed t-tests were conducted using the number of images (n = 25 for each category) as the sample size. Categorization accuracy for all categories except bedroom (PS, SS3), highway (SS3), living room (SS3), and office (PS, SS3) was significantly higher than chance level (10 %) (*p* < 0.05, FDR corrected, |*t*(24)| ≥ 2.82).

**Figure 4.**
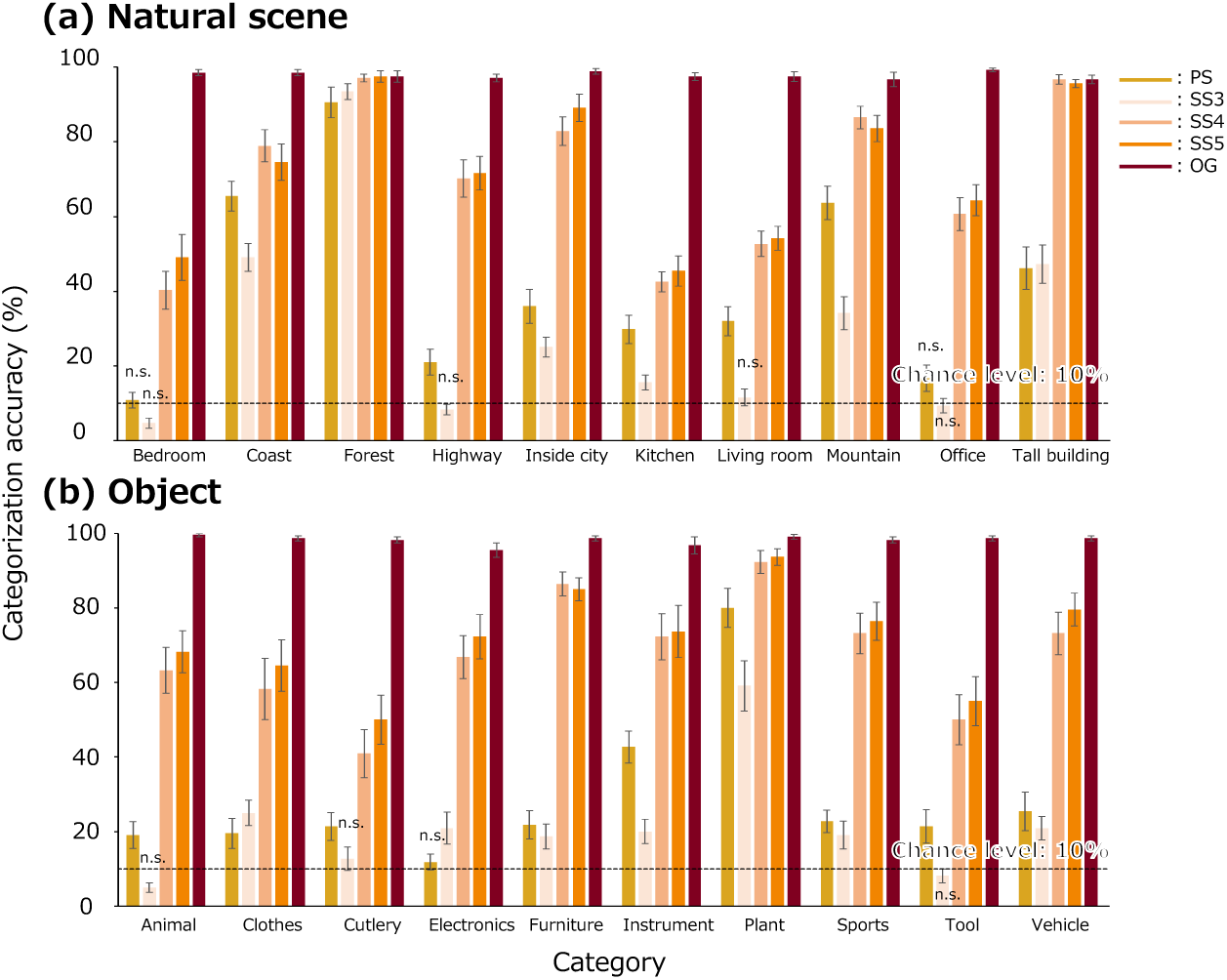
(a) Mean categorization accuracy for each scene category. Yellow bars indicate PS-synthesized images (PS), orange bars represent the three types of style-synthesized images (SS3, SS4, SS5), and red bars indicate original images (OG). The dotted line shows the chance level (10 %) for the task. Error bars represent ±1 standard error of the mean (SEM) across images. (b) Mean categorization accuracy for each object category. Yellow bars correspond to PS-synthesized images (PS), orange bars correspond to style-synthesized images (from left to right: SS3, SS4, SS5), and red bars correspond to original images (OG). The dotted line indicates the chance level (10 %). Error bars represent ±1 SEM across images. Note that all categorization accuracies exceeded the chance level significantly (p < 0.05, FDR corrected), except for conditions marked as “n.s.” (not significant).

Figure 4b shows the mean accuracy of object categorization for each category. Similar to the results for natural scene images, accuracy for OG images was consistently high across all categories. For synthesized images, the overall trend was comparable across categories, although some category-specific differences were observed. For instance, the plant category showed relatively high categorization accuracy even in the PS and SS3 conditions. Two-tailed t-tests (n = 20 for each category) revealed that categorization accuracy for all categories, except animal (SS3), cutlery (SS3), electronics (PS), tool (SS3), and vehicle (SS3), was significantly higher than chance level (*p* < 0.05, FDR corrected, |*t*(19)| ≥ 2.37).

#### Distribution of behavioral responses

Differences in categorization accuracy across categories may be attributed to biases in confusion patterns between specific category pairs. To examine response tendencies among observers, we constructed confusion matrices of category responses for each image type (PS, SS3, SS4, SS5, OG), and applied t-distributed stochastic neighbor embedding (t-SNE) to compress the response frequency data into two-dimensional space. The random seed for t-SNE was fixed to ensure the reproducibility of the results. The resulting low-dimensional embeddings were plotted to visualize the distribution of observer responses (Figure 5). As shown in Figure 4, response patterns for PS and SS3 were similar to each other, as were those for SS4 and SS5. Therefore, for simplicity, we focused on the PS and SS5 conditions.

**Figure 5.**
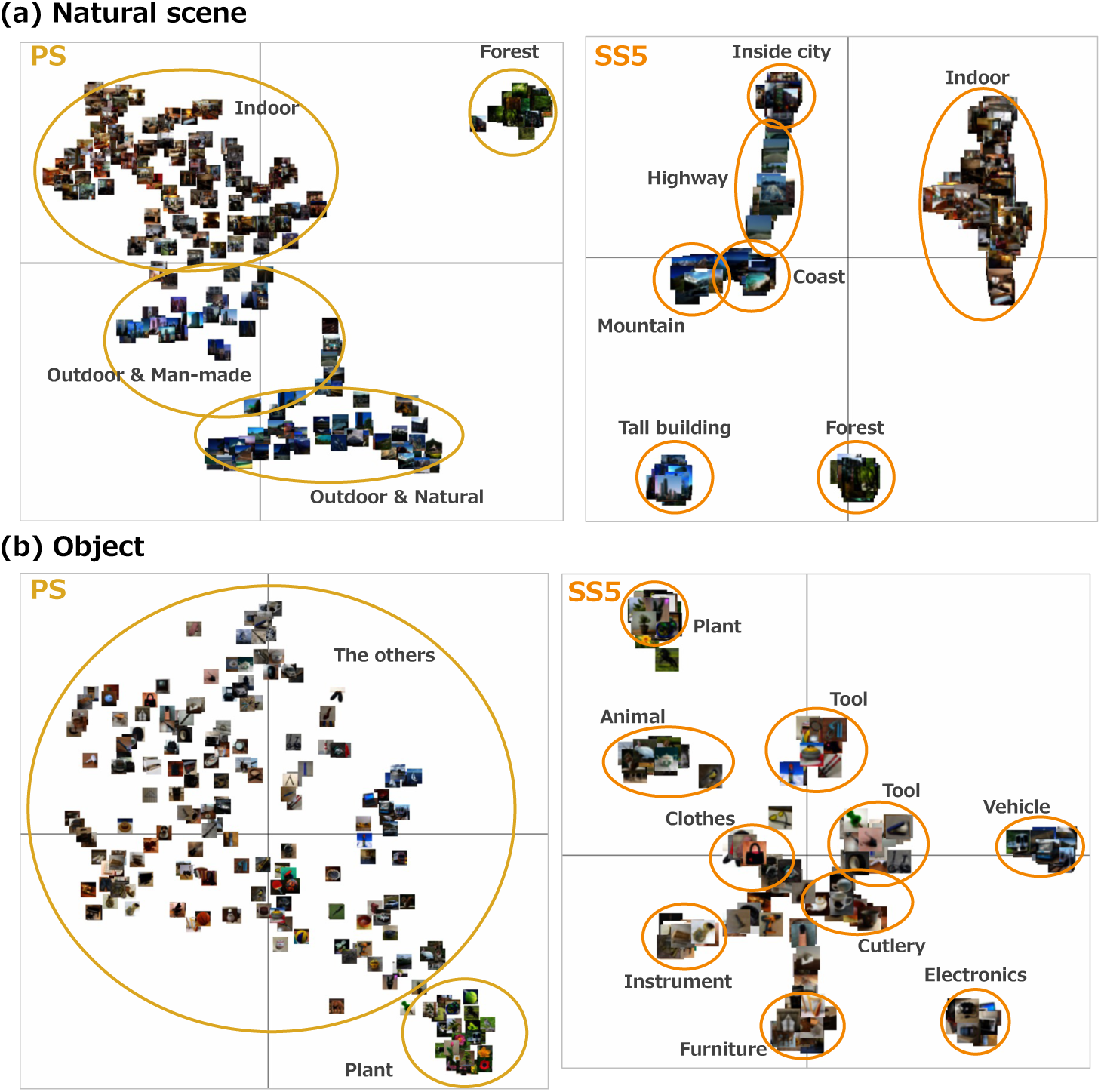
(a) Two-dimensional visualization of the observers’ natural scene categorization responses created using t-distributed stochastic neighbor embedding (t-SNE). The cross lines indicate the two coordinate axes. Labels were added to indicate regions where images densely clustered. (b) Two-dimensional visualization of observers’ object categorization responses, created using the same procedure as in (a). We note that images were blurred to address copyright issues.

Figure 5a shows the visualization of natural scene categorization. Under the PS condition, although some distinct clusters were observed for categories such as forest and mountain, the overall structure primarily separated open versus closed scenes. In contrast, under the SS5 condition, most natural scene categories, with the exception of indoor scenes such as living room, kitchen, and bedroom, formed clearly distinguishable clusters. These results suggest that under SS5, natural scene categories were generally well discriminated, except for indoor categories.

Figure 5b shows the visualization of object categorization. In the PS condition, only the plant category formed a cluster, while the remaining categories were broadly dispersed. In contrast, under the SS5 condition, nearly all object categories formed specific clusters, indicating that a majority of object images were accurately and consistently identified even for style-synthesized images.

## 2: Natural scene and object category classification from VEPs

The results of the psychophysical experiments demonstrated that observers were able to accurately categorize natural scene and object images into specific categories even when viewing style-synthesized images. However, since the presentation duration in the psychophysical experiment was fixed at 200 ms, these results do not provide insights into the temporal dynamics of how statistical image features are utilized during natural scene and object recognition. To further investigate the temporal mechanisms underlying natural scene and object recognition, we conducted an EEG experiment. In this experiment, we recorded observers’ brain activity while they viewed a subset of the visual stimuli used in the psychophysical experiment: PS-synthesized images (PS), style-synthesized images with features from all convolutional blocks (SS5), and original images (OG). We analyzed the visual evoked potentials (VEPs) to explore the time course of neural processes involved in recognizing natural scenes and objects.

Among the three types of style-synthesized images (SS3, SS4, SS5) used in the psychophysical experiment, only SS5 images were employed in the EEG experiment. This decision was based on the results of our behavioral experiment, which indicated that SS5 images sufficiently captured the contribution of style features to recognition performance. Furthermore, using only SS5 images reduced the experimental burden on observers during EEG recording.

### Materials & Methods

#### Apparatus

Visual stimuli were generated using a PC (HP Z2 Workstation) and presented on an LCD monitor (BENQ XL2730-B). The monitor was gamma-corrected (ColorCal II, Cambridge Research Systems), and the refresh rate was set to 60 Hz. The mean background luminance of the monitor was 47 cd/m². EEG signals were recorded using an EEG system (BrainVision Recorder) with a BrainAmp amplifier and EasyCap electrode cap (Brain Products GmbH).

#### Observers

In the EEG experiment using natural scene images, a total of 15 observers took part in the EEG recordings, including the three authors, as well as 12 naïve students (age range: 21–30 years, mean age: 23.9 years). All observers had normal or corrected-to-normal vision. The experiment was approved by the “Ethical Review Committee for Experimental Research on Human Subjects, Faculty of Arts and Sciences, The University of Tokyo.” Written informed consent was obtained from all observers prior to participation. Among these 15 observers, 11 observers (the three authors, and eight naïve students) had participated in the previous psychophysical experiment before taking part in the EEG experiment.

In the EEG experiment using object images, a separate group of 15 observers (including the authors) took part in the study (age range: 21–30 years, mean age: 23.8 years), following the same ethical procedures and experimental guidelines as in the natural scene image experiment.

#### Visual stimulus

For natural scene images, a total of 750 images were used in the EEG experiment. These included 250 original images (OG; 5.7 deg × 5.7 deg), 250 PS-synthesized images (PS) generated from the original images, and 250 style-synthesized images using five convolutional layers (SS5; hereafter referred to as SS).

Similarly, for object images, a total of 600 images were used, consisting of 200 original images (OG), 200 PS-synthesized images (PS), and 200 style-synthesized images (SS5; hereafter referred to as SS), all generated following the same procedures as those for the natural scene images.

#### EEG recordings and preprocessing

EEG signals were recorded from 31 electrodes (Fp1, Fp2, F3, F4, C3, C4, P3, P4, O1, O2, F7, F8, T7, T8, P7, P8, Fz, Cz, Pz, FC1, FC2, CP1, CP2, FC5, FC6, CP5, CP6, FT9, FT10, TP9, and TP10) placed according to the international 10–20 system. The signals were sampled at 1000 Hz using BrainVision Recorder, BrainAmp amplifier, and EasyCap (Brain Products GmbH). Electrode impedance was kept below 5 kΩ whenever possible. In cases where impedance could not be reduced sufficiently despite careful preparation, the experiment was conducted with the lowest impedance achievable to minimize observers’ burden. An additional electrode positioned between Fz and AFz was used as the ground. All electrodes were initially referenced online to an electrode placed between Fz and Cz. Subsequently, EEG data were re-referenced offline to the average of all electrodes. The recorded EEG data were band-pass filtered between 0.5 Hz and 40 Hz. The continuous data were then segmented into epochs from -400 ms to 800 ms relative to stimulus onset for each trial. Baseline correction was applied using the EEG signal from -100 ms to 0 ms prior to stimulus onset. Artifacts caused by eye movements were removed using independent component analysis (ICA). Additionally, any epochs containing abnormal potentials exceeding ±75 µV were excluded from further analysis. The preprocessing procedures followed the settings and methods used in our previous study (Orima & Motoyoshi, 2023).

#### Experimental design and statistical analyses

The EEG experiment was conducted in a shielded dark room. EEG recording was performed in three separate blocks, each corresponding to one of the three image types: PS-synthesized images (PS), style-synthesized images (SS), and original images (OG). For natural scene images, each block consisted of five sessions, and in each session, 250 natural scene images corresponding to that block were presented in a random order. In each trial, a visual stimulus was presented for 500 ms at the fixation point at the center of the display. Observers were instructed to maintain their gaze on the fixation point throughout the experiment. To prevent recorded EEG signals from being affected by the other tasks such as fixation task than observation of visual stimulus per se (Braun & Sagi, 1990; Lee et al., 1999; Motoyoshi, 2011; Motoyoshi et al., 2015), observers just viewed visual stimulus passively during the experiment. Observers were monitored at the end of each run and confirmed that they remained focused on the experiment. The inter-stimulus interval (ISI) was set to 750 ms (cf. Orima & Motoyoshi, 2021). The order of the three blocks (PS, SS, OG) was counterbalanced across observers. For object images (200 images × 3 conditions), EEG recordings were conducted on a different day from the natural scene image sessions, following the same procedure as described above.

To understand the temporal mechanisms underlying natural scene and object categorization, we conducted decoding analyses using a support vector machine (SVM) classifier. Specifically, we aimed to predict the natural scene or object category corresponding to each VEP. Since each image was presented five times per observer, the VEPs associated with a given image remained relatively noisy even after the preprocessing. Therefore, to improve the signal-to-noise ratio while setting the number of images as the sample size to preserve generalizability across images, VEPs were averaged not only within observers but also across observers, resulting in a single representative VEP for each image (Orima & Motoyoshi, 2021; Orima et al., 2025). For the decoding analysis, we used EEG signals recorded from 29 electrodes (F3, F4, C3, C4, P3, P4, O1, O2, F7, F8, T7, T8, P7, P8, Fz, Cz, Pz, FC1, FC2, CP1, CP2, FC5, FC6, CP5, CP6, FT9, FT10, TP9, TP10), based on the international 10–20 system. For each classification, the EEG data from all electrodes were concatenated (flattened), resulting in a feature vector with 29 × T dimensions for each image (where T is the number of time points). To examine both the contribution of each time window to classification performance and the cumulative contribution of the patterns up to each time point, we conducted two types of analyses: one based on individual time windows and the other based on cumulative data. In the window-based analysis, we used 50 ms time windows with a step size of 25 ms (i.e., T = 1–50, 26–75, …, 451–500 ms). In contrast, in the cumulative-time analysis, data were incrementally accumulated by adding 25 ms time periods starting from the initial time-window (i.e., T = 1–50, 1–75, …, 1–500 ms). We note that we applied principal component analysis (PCA) (c.f. Orima et al., 2025) in the cumulative-time analysis to reduce the feature dimensionality of each sample of the input data for the SVM model. Principal components were selected until the cumulative explained variance reached 99 %. The number of selected components was 51-202 for the natural scene classification task and 48-165 for the object classification task, both of which were smaller than the number of samples (images). Multi-class classification was performed using the one-vs-one approach, implemented via the ʻfitcecoc’ function in MATLAB. To ensure that no information leaked between training and testing data, PCA was performed independently within each cross-validation fold.

The significance of analyzed data was tested by binomial test using the number of images (250 for natural scene, 200 for object) as the sample size. We applied the false discovery rate (FDR) correction method proposed by Benjamini & Hochberg (1995) to the p-values calculated at each data point, to address the multiple comparisons.

### Results

#### Visual evoked potentials (VEPs)

Figure 6 shows representative visual evoked potentials (VEPs) recorded from typical electrodes (O1/O2, Pz, and F3/F4) in response to PS-synthesized images (PS), style-synthesized images (SS), and original images (OG). The time course of the VEPs is shown for each condition. The topographical maps represent scalp distributions of the VEPs, where red indicates positive potentials and blue indicates negative potentials.

**Figure 6.**
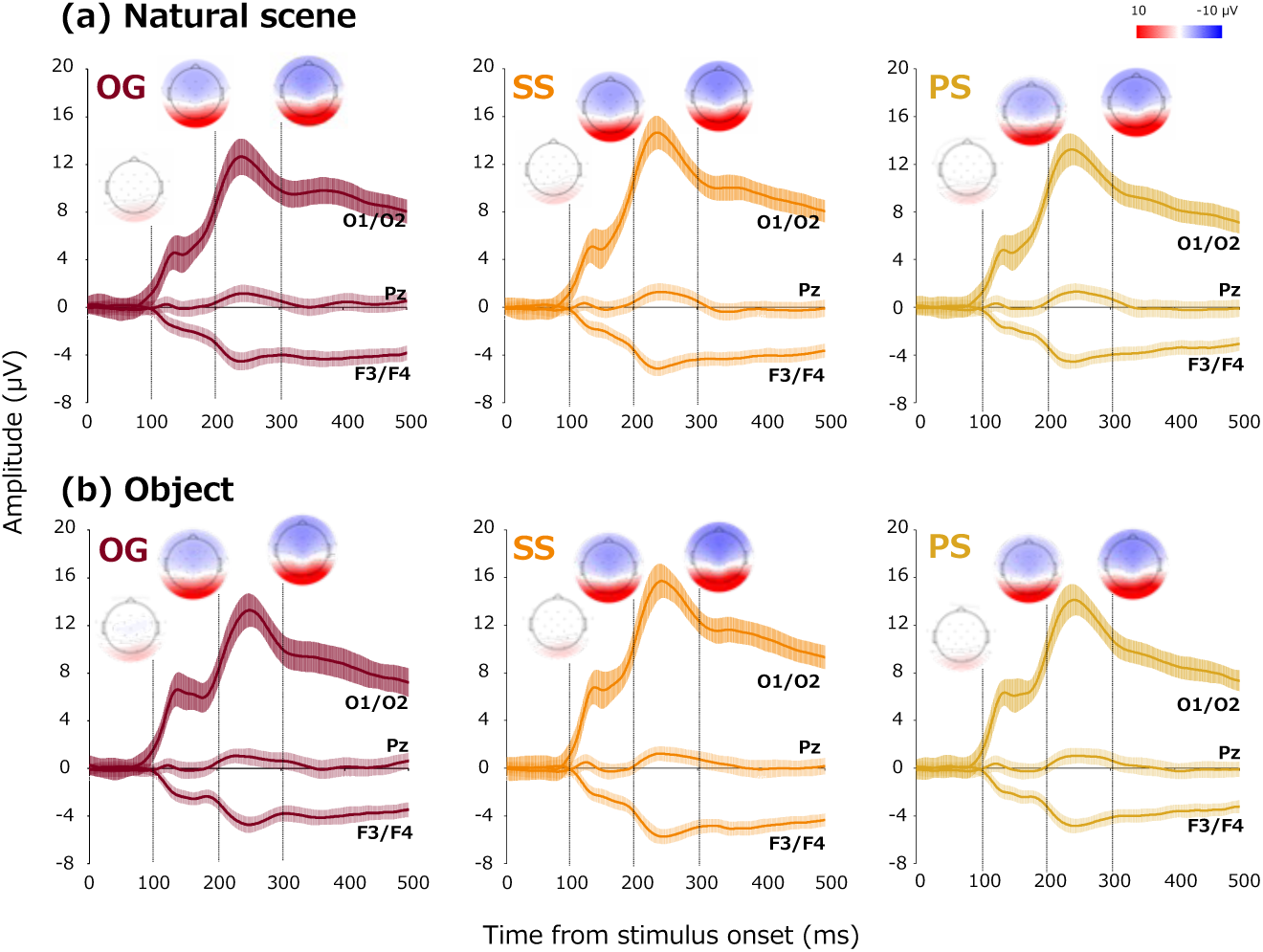
(a) Visual evoked potentials (VEPs) recorded while observers viewed natural scene images. Solid lines represent the grand average across observers for each electrode, and shaded areas indicate ±1 standard deviation (SD) across images. The time points for the topographical maps correspond to the vertical dashed lines. (b) Visual evoked potentials recorded while observers viewed object images. Topographical maps are shown using the same method as in (a).

#### Natural scene and object category classification from VEPs

Figure 7a shows the natural scene classification accuracy over time achieved by SVMs trained on VEPs for PS-synthesized (PS), style-synthesized (SS), and original (OG) images. Using the number of original images (n = 250) as the sample size, we conducted binomial tests at each time point. The results of the cumulative-time analysis indicated that classification performance significantly exceeded the chance level (10 %) from 175 ms (OG) and 150 ms (PS and SS) onward (*p* < 0.05, FDR-corrected). The peak classification accuracies were 25.6 % for PS, 32.0 % for SS, and 34.4 % for OG. The time-window based analysis showed that the classification accuracy reached the peak at from 150-200 ms to 251-300 ms time-window.

**Figure 7.**
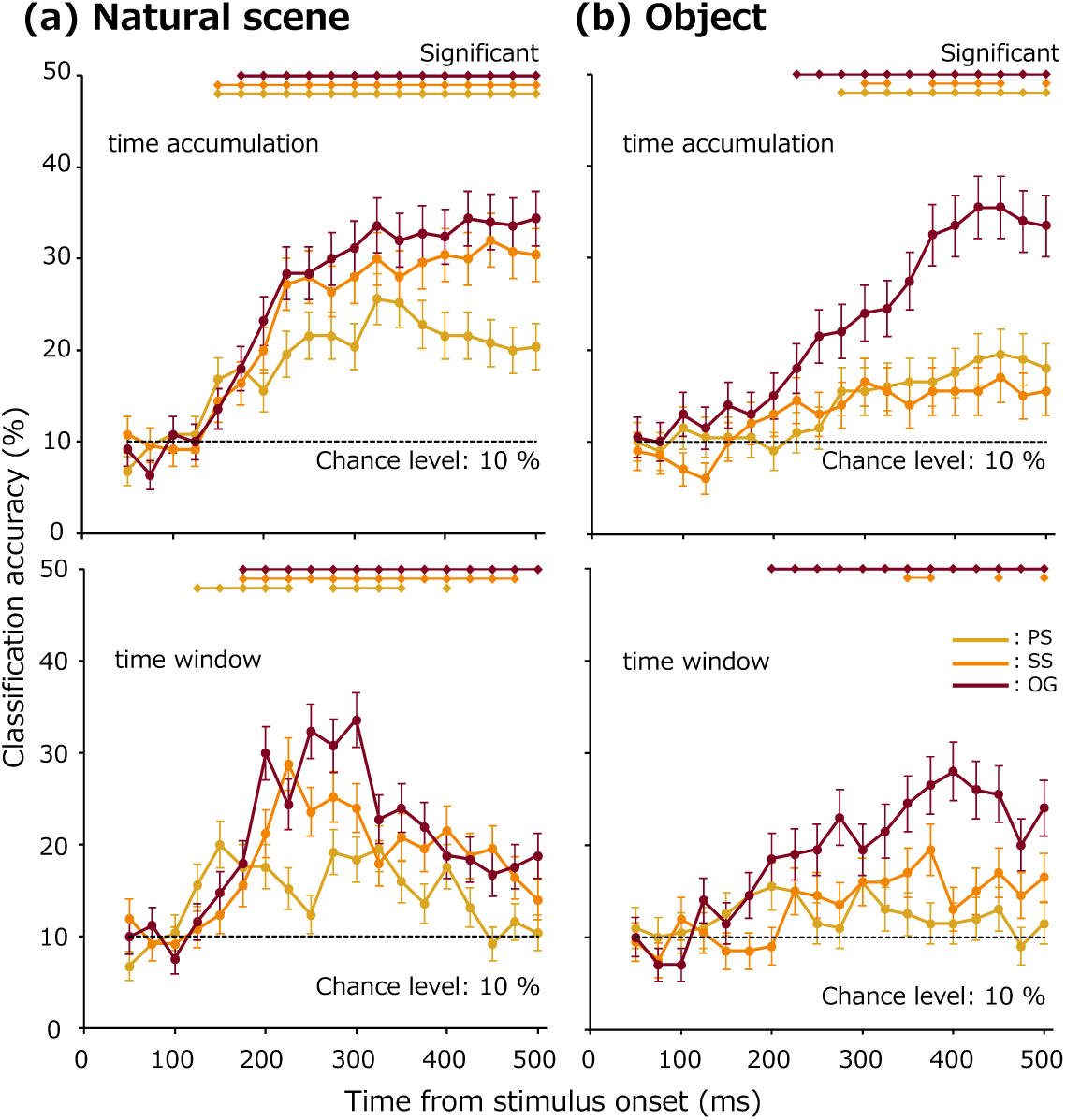
(a) Time course of natural scene category classification accuracy based on SVM models trained on VEPs. Classification accuracy was calculated as the average across 250 individual models, each trained on a different training set. The horizontal axis indicates the end period of the VEP time window used for training the model. The yellow line corresponds to models trained on VEPs for PS-synthesized images (PS), the orange line to those for style-synthesized images (SS), and the red line to those for original images (OG). Dotted segments along the top of the graph indicate time points where classification accuracy was significantly above chance level (p < 0.05, FDR-corrected). (b) Time course of object category classification accuracy from VEPs, plotted using the same method as in (a).

Figure 7b shows the corresponding results for object category classification. Using the number of original images (n = 200) as the sample size, binomial tests for the cumulative-time analysis revealed that classification accuracy was significantly above the chance level from 225 ms (OG), 300 ms (SS), and 275 ms (PS) onward (*p* < 0.05, FDR-corrected). The peak classification accuracies were 19.5 % for PS, 17.0 % for SS, and 35.5 %for OG. The time-window based analysis showed that the classification accuracy reached the peak at 351-400 ms time-window.

These results indicate that natural scene categories could be classified with relatively high accuracy based on VEPs for original images (OG) or style-synthesized images (SS), which suggests that information reflected in the VEPs recorded while viewing OG or SS images contributed substantially to natural scene category classification.

In contrast, for object categories, high classification accuracy was achieved only when VEPs for OG images were used. In addition, the latency that this classification became significant was later than that of natural scene images. Furthermore, classification accuracy based on VEPs for SS or PS images was considerably lower. These results suggest that while VEPs for OG contained sufficient information for significant object category classification, VEPs for SS contained information sufficient for statistically significant, but not highly accurate, classification of object categories.

## 3: Contribution of style features to natural scene and object category classification

Results from the psychophysical experiment and VEP-based analyses indicated that style-synthesized images (SS) of both natural scenes and objects contained sufficient information for category-level discrimination. However, differences in VEP-based classification performance suggest that the human visual system may utilize different types of information for natural scene or object recognition. To directly evaluate the contribution of style features to category discrimination, we conducted an analysis to determine how accurately the style features themselves, which were computed from each visual stimulus, could classify natural scene and object categories.

### 3-1. Category classification based on style features

#### Materials & Methods

##### Experimental design and statistical analyses

Style features were extracted from the original images (OG) and PS-synthesized images (PS), using the same procedure as in neural style transfer (NST). Since the style features extracted from OG images are theoretically equivalent to those preserved in the corresponding style-synthesized (SS) images, style features were not extracted directly from the SS images. The extracted style features consisted of approximately 600,000 dimensions per image. To reduce dimensionality, principal component analysis (PCA) was conducted separately for each of the following four conditions: OG (scene), PS (scene), OG (object), and PS (object). Principal components were retained until the cumulative explained variance exceeded 99 %. The number of retained components was 74-76, 61-62, 89-90, and 72-74, respectively, for each of the four conditions. To ensure that no information leaked between training and testing data, PCA was performed independently within each cross-validation fold.

For classification, we performed a leave-one-out cross-validation separately for each condition (OG and PS x natural scene and object). In each fold, the style feature computed from one image was assigned as the test sample, while those from the remaining images served as the training set for an SVM classifier for predicting the correct category. By comparing the predicted category with the ground truth, classification accuracy was calculated for each image. To simulate the variation across observers in the psychophysical and EEG experiments, we incorporated variability into the training data. Concretely, for each fold, the training set consisted of the style features computed from n/2 images randomly selected from the remaining (n-1) images (excluding the test image), and this sampling process was repeated 15 times with minimally overlapping subsets. An SVM classifier was trained on each of these 15 subsets, and predictions were made for the testing image. Final classification accuracy for each image was computed as the average accuracy across the 15 independently trained classifiers. The number of repetitions (15) was chosen to match the number of observers in the EEG experiment.

The statistical significance of analyzed data was tested by permutation test using the number of images (250 for natural scene, 200 for object) as the sample size. We applied the false discovery rate (FDR) correction method proposed by Benjamini & Hochberg (1995) to the p-values calculated at each data point, to address the multiple comparisons.

#### Results

Figure 8a shows the classification accuracy for natural scene categories, and Figure 8b shows the classification accuracy for object categories. In both figures, the average classification accuracy across all categories and the accuracy for each individual category are shown. The average classification accuracy was 76.1 % (scene, OG), 53.9 % (scene, PS), 52.2 % (object, OG), and 33.6 % (object, PS). We conducted permutation tests (10,000 times iteration) using the number of images (250 for natural scene, 200 for object) as the sample size, comparing classification accuracy against the chance level (10 %). The results indicated that all classification accuracies were significantly above chance (*p* ∼ 0, for average across natural scene categories; *p* ∼ 0, for average across object categories; *p <=* 10^-4^ for each natural scene category, and *p* <= 0.0321 for each object category; thresholding *p* < 0.05, FDR-corrected), except for the accuracy for clothes (PS) in the object category. Style features extracted from OG natural scene images enabled highly accurate classification of natural scene categories, achieving approximately 80 % accuracy. In contrast, style features extracted from OG object images yielded a lower accuracy of approximately 55 % for object category classification. When comparing category-wise accuracy patterns with those obtained from the behavioral experiments (Figure 4), we found that they are consistent with behavioral data for natural scene categories, but not for object categories.

**Figure 8.**
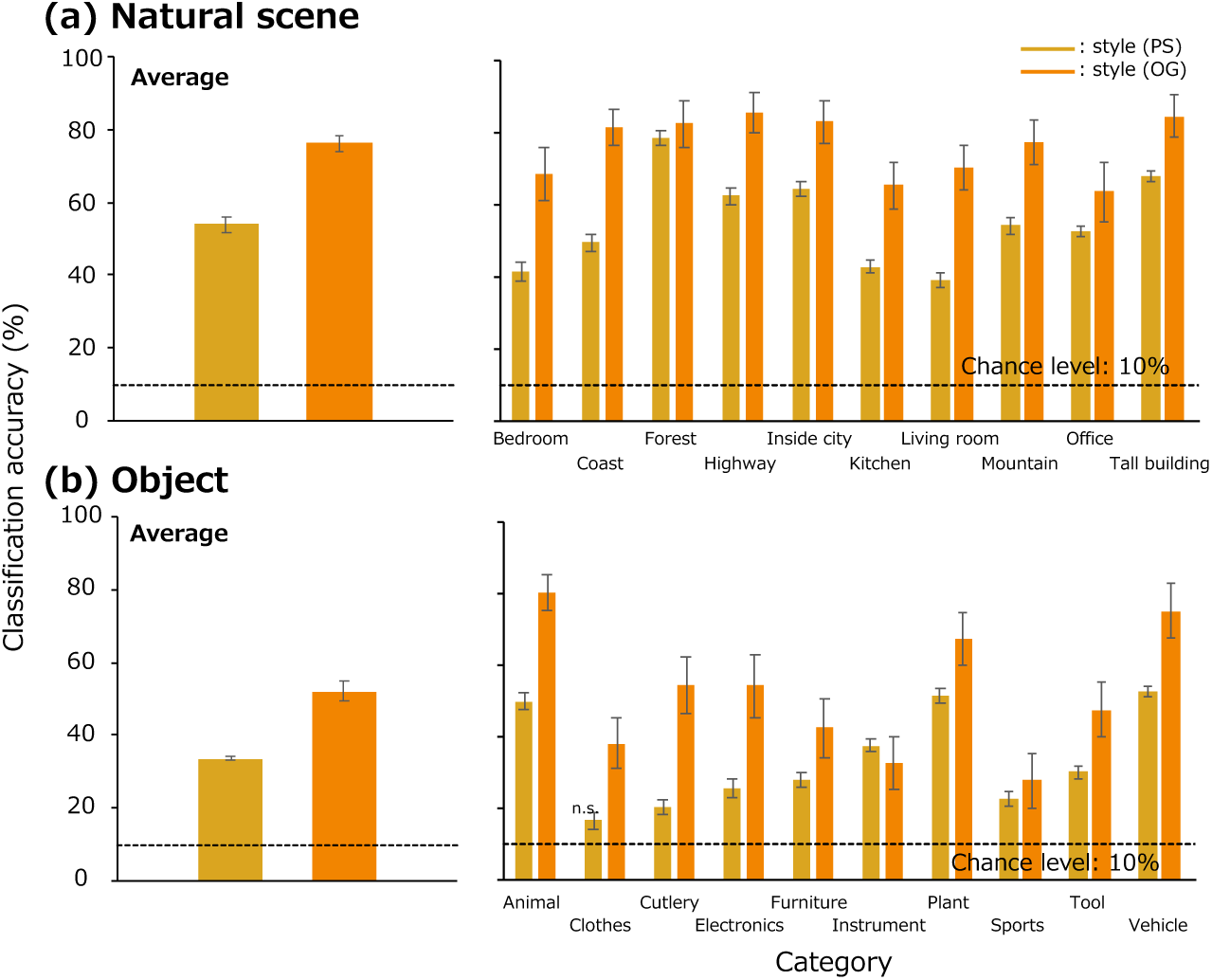
(a) Classification accuracy of natural scene categories based on compressed representations of style features extracted from natural scene images (OG, PS). The figure shows both the average accuracy across all categories and the accuracy for each individual category. (b) Classification accuracy of object categories based on compressed representations of style features extracted from original object images. The format is the same as in (a). Classification accuracies were significantly higher than the chance level (p < 0.05, FDR-corrected) for all categories except those marked as “n.s.” (not significant) in the figure.

To quantitatively examine this consistency, we computed Pearson’s correlation coefficients (*N* = 10 categories) between the behavioral accuracy and the style-feature-based classification accuracy for each category. The results are shown in Figure 9. For natural scene categories based on OG images, classification accuracy from style features was strongly correlated with observers’ behavioral performance using SS images (*r* = 0.81, *p* < 0.005). In contrast, for object categories based on OG images, the correlation was weak and not significant (*r* = 0.13, *p* > 0.7). For PS images, the correlation between style-feature-based classification and behavioral performance was not statistically significant for both natural scene categories (*r* = 0.55, *p* = 0.093), and object categories (*r* = 0.53, *p* > 0.1). These results suggest that style features extracted from OG natural scene images allow for highly accurate prediction of natural scene categories, and the prediction pattern closely matches observers’ behavioral data. In contrast, these findings were not true for object images, supporting our previous assumption that the contribution of style features differs between natural scenes and objects.

**Figure 9.**
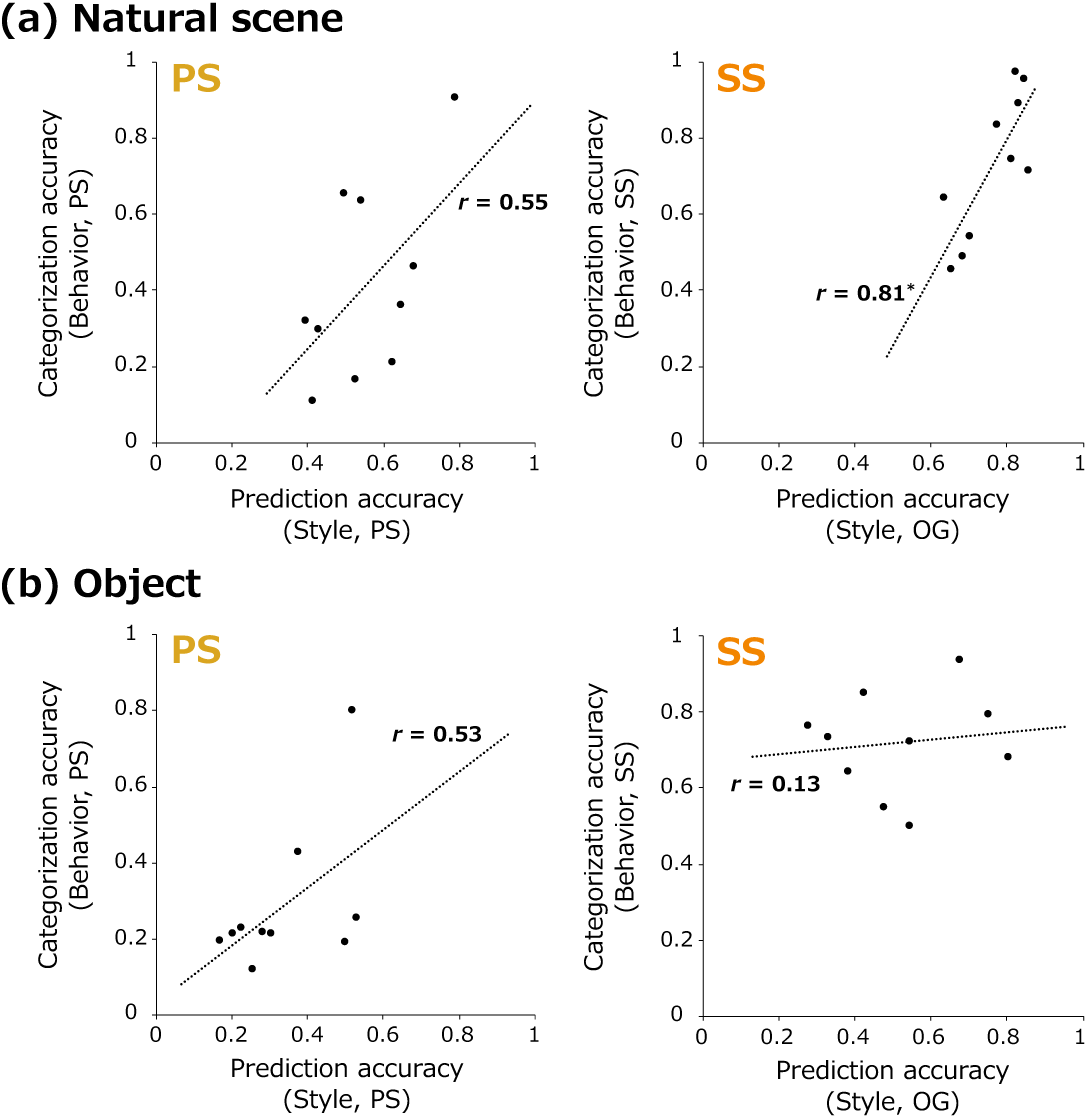
(a) Scatter plots showing the relationship between the category-wise classification accuracy based on style features extracted from original (OG) and PS-synthesized (PS) natural scene images, and the categorization accuracy of observers for each natural scene category (from SS and PS images, respectively). (b) Scatter plots showing the relationship between the category-wise classification accuracy based on style features extracted from original (OG) and PS-synthesized (PS) object images, and the categorization accuracy of observers for each object category (from SS and PS images, respectively). In both plots, asterisks (*) indicate that the correlation coefficient was statistically significant (p < 0.05).

### 3-2. Analysis of the encoding dynamics of style features

To investigate how style features contained in natural images are temporally encoded in the human visual cortex, we conducted a reverse correlation analysis between visual evoked potentials (VEPs) for natural scene and object images and the corresponding style features.

#### Materials & Methods

##### Experimental design and statistical analyses

We conducted a reverse correlation analysis to examine the relationship between visual evoked potentials (VEPs) elicited by original images (OG) of natural scenes and objects, and the style features computed from those images. VEPs were analyzed within sliding time windows of 50 ms, starting from stimulus onset: 1–50 ms, 26–75 ms, …, up to 451–500 ms. We used data from 29 electrodes excluding Fp1 and Fp2, which are susceptible to eye movement artifacts, consistent with our previous analyses. For each time window, VEP data from all time points and electrodes were flattened, resulting in 29 × 50 variables per image. To reduce the dimensionality of the VEP data, principal component analysis (PCA) was applied (c.f. Orima et al., 2025), and principal components were retained until the cumulative explained variance reached 99 %. The number of retained components ranged from 58 to 79 across time windows.

Similarly, PCA was applied to the style features extracted from each image, which originally contained over 610,000 dimensions, to reduce dimension. The number of principal components was determined so that the cumulative explained variance exceeded 99 %, resulting in 196 components for natural scenes and 176 components for objects. To account for the difference in the number of variables between VEPs and style features, we computed dissimilarity matrices separately for the compressed VEP data (within each time window) and the compressed style features. Finally, we calculated the correlation coefficients between these two distance matrices (natural scenes: *N* = 31,125 pairs; objects: *N* = 19,900 pairs). We applied the false discovery rate (FDR) correction method proposed by Benjamini & Hochberg (1995) to the p-values calculated at each data point, to address the multiple comparisons.

#### Results

Figure 10 shows the temporal dynamics of the correlation between dissimilarity matrices computed from compressed VEP data and compressed style features. To enable a fair comparison between the two conditions with different sample sizes (natural scenes vs. objects), we plotted the absolute t-values instead of correlation coefficients.

**Figure 10.**
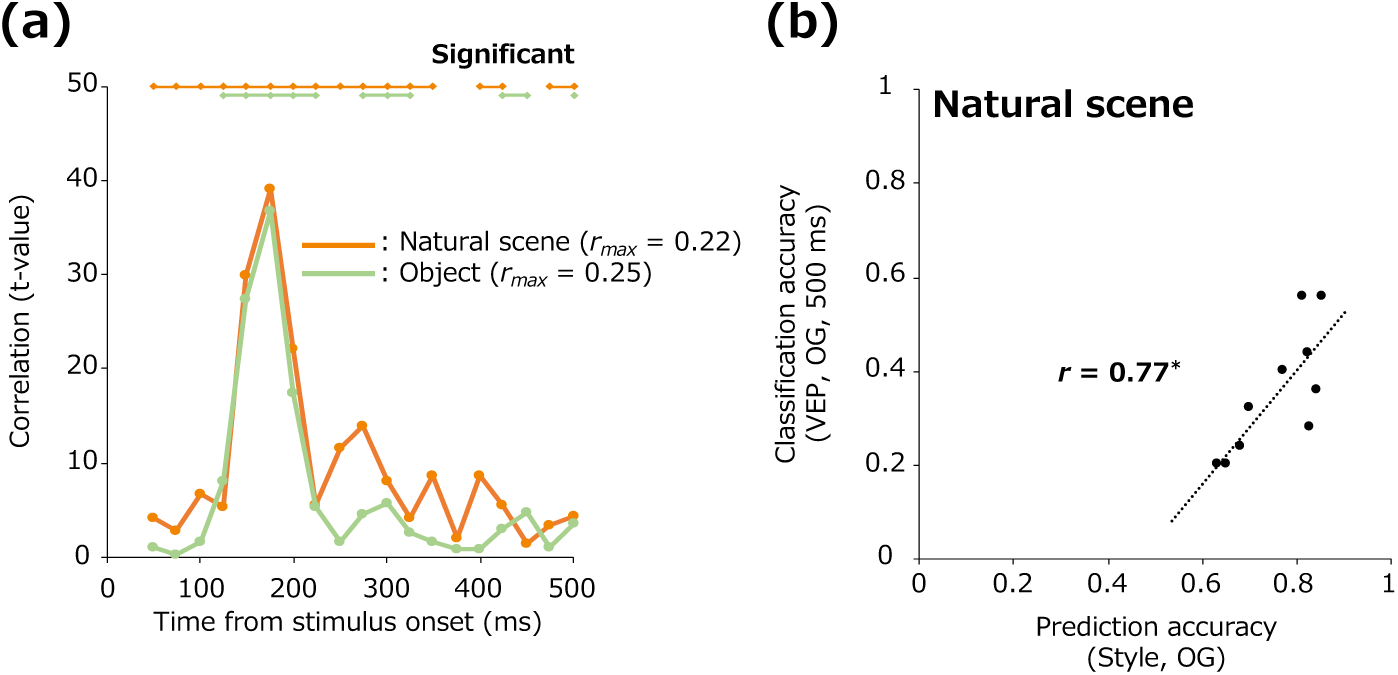
(a) Temporal dynamics of the correlation between VEPs for natural scene or object images (OG) and the style features extracted from the corresponding images. The horizontal axis indicates time from stimulus onset (ms), and the vertical axis indicates the t-values of the correlation. Horizontal lines above the plots indicate time windows where the correlation was statistically significant (p < 0.05, FDR-corrected). (b) Scatter plot showing the correlation between natural scene category-wise classification patterns based on VEPs (recorded within 500 ms after stimulus onset) and those based on style features extracted from OG natural scene images. An asterisk (*) indicates that the correlation coefficient was statistically significant (p < 0.05).

For both natural scene and object images, the highest correlation was observed for VEPs in the 150–200 ms time window after stimulus onset, and this correlation was statistically significant. Notably, this latency was consistent with that reported in the previous study (Orima et al., 2025), suggesting that the compressed representation of style features is encoded at a relatively early stage of visual processing, supporting the perception of both natural scene and object images. Additionally, following the same procedure described in the previous section, we examined the correlation between natural scene category-wise classification accuracy based on VEPs and that based on style features extracted from OG natural scene images. A strong and significant correlation was observed (*r* = 0.77, *p* < 0.01), further supporting the idea that category information reflected in VEPs for natural scene images is closely related to statistical image features captured by style features.

## Discussion

In the present study, we investigated the role of statistical image features in natural scene and object recognition. To this end, we conducted natural scene and object category classification experiments using synthesized images that shared specific statistical image features with the original images (OG): PS-synthesized images (PS), which preserved PS statistics (lower-level statistical image features), and style-synthesized images (SS), which preserved style features (higher-level statistical image features). The behavioral results showed that when observers viewed style-synthesized images generated using a larger number of convolutional layers (SS4 and SS5), they were able to categorize natural scenes and objects with an accuracy of 70 % or more. This finding suggests that statistical image features play a crucial role in the recognition of both natural scene and object category. However, VEP-based decoding results revealed a difference between natural scenes and objects. While VEPs for SS images allowed for accurate classification of natural scene categories, classification accuracy for object categories remained low. These results imply that while natural scene categories are perceived based on the statistical image features that are both present in SS images and reflected in neural responses, object categories are likely perceived based on features that are present in SS images but not strongly represented in VEPs. To further examine this possibility, we directly tested whether style features themselves could classify natural scene and object categories. The results showed that style features enabled highly accurate classification of natural scene categories but only moderate classification accuracy for object categories. Moreover, the category-wise accuracy pattern for natural scenes was consistent with observers’ behavioral data and VEP-based decoding, whereas the pattern for objects was inconsistent with these measures. Taken together, these findings support the idea that the perception of natural scene categories is largely supported by statistical image features such as style features, whereas the contribution of style features to object category recognition remains unclear.

As shown in Figures 3 and 4, style-synthesized images contained sufficient information to enable highly accurate classification of natural scene categories. Furthermore, as demonstrated in Figure 7a, VEPs for original natural scene images or style-synthesized images allowed for statistically significant natural scene category classification from approximately 175 ms after the stimulus onset. This finding suggests that neural responses evoked by both original and style-synthesized images reflected information sufficient for natural scene category classification. Therefore, the high classification accuracy of natural scene categories based on VEPs elicited by both original and style-synthesized images suggests that the features contributing to natural scene categorization were likely global statistical features, namely style features, which were shared across these images. This interpretation is not inconsistent with the fact that EEG inherently captures spatially summed activity of large populations of neurons (Nunez & Srinivasan, 2006), which enabled previous studies to demonstrate that EEG signals encode the global impression or statistical properties of visual images (Scholte et al., 2009; Groen et al., 2013; Greene & Hansen, 2020; Orima & Motoyoshi, 2021; Orima et al., 2025).

In the decoding of natural scene and object categories from VEP data, the time-window based analysis revealed potential differences in the hierarchical processing of these categories in the brain. The classification accuracy for natural scene categories first reached statistical significance in 126–175 ms time window after the stimulus onset, whereas for object categories, significant accuracy was firstly observed 151–200 ms time-window. These findings are consistent with previous EEG and MEG studies suggesting that the perception of scene categories and the peak decoding accuracy for faces and objects typically emerge around 150–200 ms after the stimulus onset (Harel et al., 2016; Mohsenzadeh et al., 2017; Lowe et al., 2018; Kaiser et al., 2020). However, while the peak classification accuracy for natural scene categories appeared at a latency of approximately 200–300 ms after the stimulus onset, the peak for object categories occurred much later, around 400 ms. This difference in peak latency may reflect distinct neural processes involved in the processing of natural scenes and object categories. Notably, the peak latency for natural scene category decoding is consistent with the latency associated with spatial layout processing (Cichy et al., 2017). On the other hand, the peak latency observed for object category decoding aligned with the timing associated with higher-level visual areas, such as the inferior-temporal cortex, rather than early visual cortices, according to comparisons between MEG and fMRI data (Cichy et al., 2014; Mohsenzadeh et al., 2017). This later latency may reflect higher-order processes such as detailed object shape analysis or feedback-related activity.

As shown in Figure 10a and proposed by a previous study (Orima et al., 2025), style features are strongly reflected in EEG signals. To directly test whether style features contributed to natural scene categorization, we examined the classification performance based solely on style features. As shown in Figure 8a, natural scene categories could be classified with an accuracy of approximately 80 %. This result suggests that the perception of natural scene categories may be robustly represented by style features. Moreover, as shown in Figure 9a, there was a very strong correlation between the category-wise classification accuracy based on style features and observers’ behavioral responses. This further supports the idea that style features significantly contribute to natural scene category recognition.

Although style features contributed substantially to natural scene category classification, they did not fully capture all aspects of natural scene category perception. The behavioral experiments demonstrated that the categorization accuracy varied depending on the natural scene category. As shown in Figure 5a, categories such as coast, forest, highway, inside city, mountain, and tall building formed distinct clusters, while indoor categories such as bedroom, kitchen, living room, and office formed a single mixed cluster. This pattern suggests that while observers were able to classify outdoor categories from style-synthesized images almost perfectly, they frequently confused indoor categories, which is consistent with a previous study (Quattoni & Torralba, 2009). These results imply that style features are highly effective for distinguishing outdoor natural scene categories but are insufficient for the classification of indoor scenes. Indeed, distinguishing between living room and kitchen likely requires more than recognizing that the scene belongs to an indoor space; it requires additional information about objects, furniture, and the spatial layout within the scene. Such processes involve more time-consuming and complex visual computations (Itti et al., 1998; Itti & Koch, 2000; Hochstein & Ahissar, 2002). Since style features essentially aggregate spatial information into global statistical features, they may fail to represent these critical local or layout-dependent details. To achieve a more precise representation of natural scene categories, a possible solution would be to compute statistical features within spatially divided blocks as proposed in previous studies (Oliva & Torralba, 2001; Lazebnik et al., 2006; Jagadeesh & Gardner, 2022), thereby preserving local spatial structure while still capturing useful statistical information.

The results for object images revealed a different pattern from those observed for natural scene images. Behavioral data suggested that style-synthesized images contained sufficient information for accurate object category recognition. However, as shown in Figure 7, object category classification based on VEPs yielded high accuracy when observers viewed original images (OG), but only marginally significant accuracy when they viewed style-synthesized images (SS). Given that object categories could be decoded from VEPs elicited by OG images, it is likely that object recognition also involves global information that is reflected in EEG signals. Indeed, recent studies have suggested that it is possible to detect animals in natural scenes (Banno & Saiki, 2015) from PS synthesized images, or decode properties such as animacy and real-world object size from EEG signals (Wang et al., 2022). The current findings suggest that statistical image features, even reflected in VEPs, may still convey meaningful information for object recognition. On the other hand, the failure to achieve high classification accuracy of object categories from VEPs for SS images suggests that the global image features, which was computed from the entire image including the background, may not have been sufficient to support object category recognition. Previous studies have indicated that object recognition relies not only on the object itself but also on contextual background information (Palmer, 1975; Bar, 2004; Torralba et al., 2006; Oliva & Torralba, 2007; Brandman & Peelen, 2017). Since the process of neural style transfer does not explicitly preserve the spatial relationships between objects and their background, it is possible that critical features for object recognition that would be reflected in VEPs were not well retained in the style-synthesized images. Nevertheless, the style-synthesized versions of these images likely contained abundant local features related to the objects themselves. Such mid-level features (Ullman et al., 2002) may have contributed to the high behavioral categorization accuracy. Taken together, these findings suggest that in both natural scene and object recognition, the recognition of specific objects placed within a scene may not be fully supported by style features alone. It highlights a potential limitation of purely statistical image features in representing fine-grained visual category information.

## Acknowledgments

The present study was supported by JSPS KAKENHI JP21J20898, JP24K2222, JP20K21803, JP23K25751, and JP24H01540.

